# A Web Service for Assessing Insect Abundances for Meadow Birds by Image Analysis

**DOI:** 10.1101/660027

**Authors:** Ricardo Michels, Rutger Vos

## Abstract

Meadow birds are a group of species native to the Netherlands characterized by breeding in meadows that has been in decline over the last several decades, despite widespread conservation efforts. Agricultural intensification is thought to be one of the main causes of this decline, but no yearly data exists on the surrounding ecology of these birds. Recent efforts have tried to assess the food supply of meadow birds by setting sticky traps and counting the number of insects caught on them. However, this approach cannot be applied on a large scale since counting the insects is very labour intensive and unappealing to the volunteers that contribute to this research. To get a better assessment of the food supply at a larger scale, we present a system to automate counting of insects on sticky traps. The system is intended to process uploaded images and metadata using computer vision techniques to determine the number of insects found in photographs taken from the sticky traps.

## Introduction

Meadow birds are a group of birds that have their primary habitat on meadows and pastures created by humans. One example of this group is the black-tailed godwit (*Limosa limosa*, in Dutch: “grutto”, Figure 1). Their eggs hatch in the first half of May (Beintema et al., 2007), and the newly hatched birds eat primarily small invertebrates (>4 mm) between 10 to 20 centimetres off the ground whilst the adults mostly feed on worms present in the soil. Populations of meadow birds in the Netherlands are threatened: their numbers have been declining steadily for a long time (Sovon, 2017). One of the possible causes for this decrease is the intensification of agricultural land use, destroying nests, killing the chicks during mowing activities, and decreasing the food supply available for the chicks, thereby lowering foraging success, as well as making the habitat unsuitable for these chicks (Schekkerman & Boele, 2008). Decreased foraging success quickly influences the growth of the chicks, and may result in increased risks of predation and disease.

**Figure 1.**
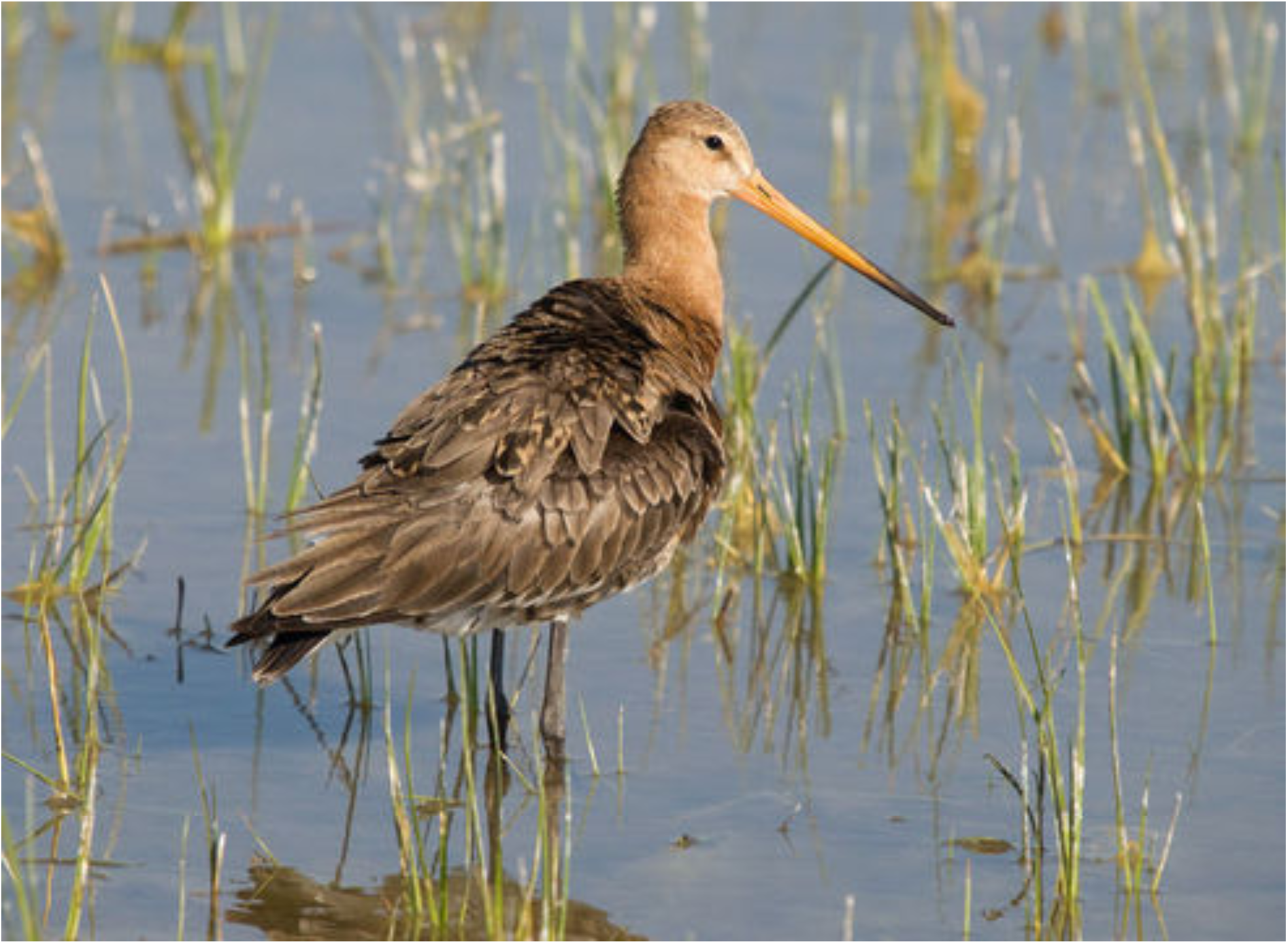
Black-tailed Godwit (Limosa limosa, L. 1758). Image from Encyclopaedia of life in public domain.

In an effort to improve the quality of the nature and ecology in the Netherlands, a system of agricultural nature management has been implemented. In this system, farmers are subsidized to make alterations to their farming practices that benefit nature. Several of these management types are aimed at helping meadow bird chicks by, for instance, safeguarding the nests using a metal framework keeping farming equipment and farm animals away, or delayed mowing, thereby giving the chicks more time to grow.

Extensive literature research did not yield any studies that systematically tried to assess the amount of food available to the meadow bird chicks during their first weeks. This information is useful in creating an effective conservation strategy and should therefore be available to policy makers considering the importance of food availability for the survival of the chicks. However, the availability of food is difficult and labour intensive to determine.

One approach to assess the amount of food available to the chicks involves counting the number of insects caught on sticky traps, with such numbers sometimes reaching well over 1000 insects per trap. Several of these traps must be deployed per field and numerous fields should be assessed to obtain a sufficiently large sample to assess the number of insects present. This is especially true if the food supply is estimated over a large area. However, counting these insects is unappealing to volunteers and is extremely labour intensive, making large scale monitoring programs impractical at the present time (Deru et al., 2016; Noordijk, 2014).

In a proposal by Deru et al. (2016) and Noordijk (2014) a monitoring program was envisioned where volunteers set traps during one weekend of the year throughout the Netherlands. To make such large-scale monitoring possible a system would have to be developed that is capable of automatically determining the number of insects present on a trap, as well as grouping the insects in different size classes. Several earlier studies have created similar programs to count the number of pests caught on sticky traps placed in greenhouses (Barbedo, 2014; Solis-Sánchez et al., 2009), but our literature research did not indicate the existence of similar programs related to conservation efforts. This is important because the difference between the present research and these earlier studies is that there are no expectations to identify the species present on these sticky traps, but that the size of the insects is important to this study, whilst size was not a factor in the identification of various pests.

This paper is connected to ongoing research into computer vision at Naturalis. One aim of this activity is to create software that can automatically identify what species an organism is based on one or more photos of it analysed using trained artificial neural networks. This has been implemented for a group of orchid species and for a large number of butterflies, while a goal at present is to determine at least the genus of several species of mosquitos using photos of their wings. This is in relation with the prevention of diseases with mosquitos as a vector. These research activities have resulted in various artefacts for deploying mobile web applications that handle image uploads as well as software libraries for managing and processing image data. These results are re-used for the present project.

## Materials and methods

The main analysis script was developed in Python 2.7.6 using the Imgpheno library developed at Naturalis^1^, and the OpenCV2 library for most computer vision functionality (OpenCV-2.4.13^2^). A website was also developed as part of this project. This website was also built in python using the Django library (Django 1.8^3^), and using the website developed as part of the OrchID website of Naturalis (Pereira et al. 2016) as a template. The google maps API^4^ and the Geodjango package (django-geoposition 0.3.0^5^) was used to provide geolocation functionality required for future research. The script is available as open source software (MIT licence) at https://github.com/naturalis/imgpheno/blob/master/examples/sticky-traps.py

### Computational workflow

The steps of the image processing system are outlined in Figure 2, and proceed as follows:

1. The images are first read into the program^6^. A colour space conversion to HSV takes place to enable the automatic detection of the traps against the background^7^. Using the HSV colour space allows an easier selection specifically for the colour yellow. A binary image is made where the yellow colour from the trap is white and the background is black^8^. This does not distinguish enough between the trap and the insects caught, so a second step thresholding operation is required for the program to operate properly.
2. The program then uses the contour of the trap as found in step 1 to find the corners of the sticky trap^9^. The image is transformed according to a perspective transformation so that it contains only the sticky trap, and all the insects present on it, with standardized dimensions. These standard dimensions are required to more easily and accurately determine the calibrated size of the insects caught.^10^
3. After the image has been transformed into standard dimensions a binary map is created separating the insects from the trap. ^11^The red colour channel has the highest contrast between the yellow trap and the insects and therefore this channel is used in the creation of the binary image. ^6^The individual insects will be segmented out by finding all individual contours, where it is assumed that one contour contains one insect. This will invariably induce some error because when insects overlap they will only be counted as one instead of two or more. This however is one of the fastest ways to segment the image, which is crucial if the project is scaled up and it should not pose a large factor if the density of the insects is low enough. ^12^
4. For each contour, the area it contains is used to determine the size of the insect and the program will output the data in an output file that can be read into a statistical program like R without requiring any further processing. This will help speed up the analysis of the data. ^13, 14^

**Fig 2.**
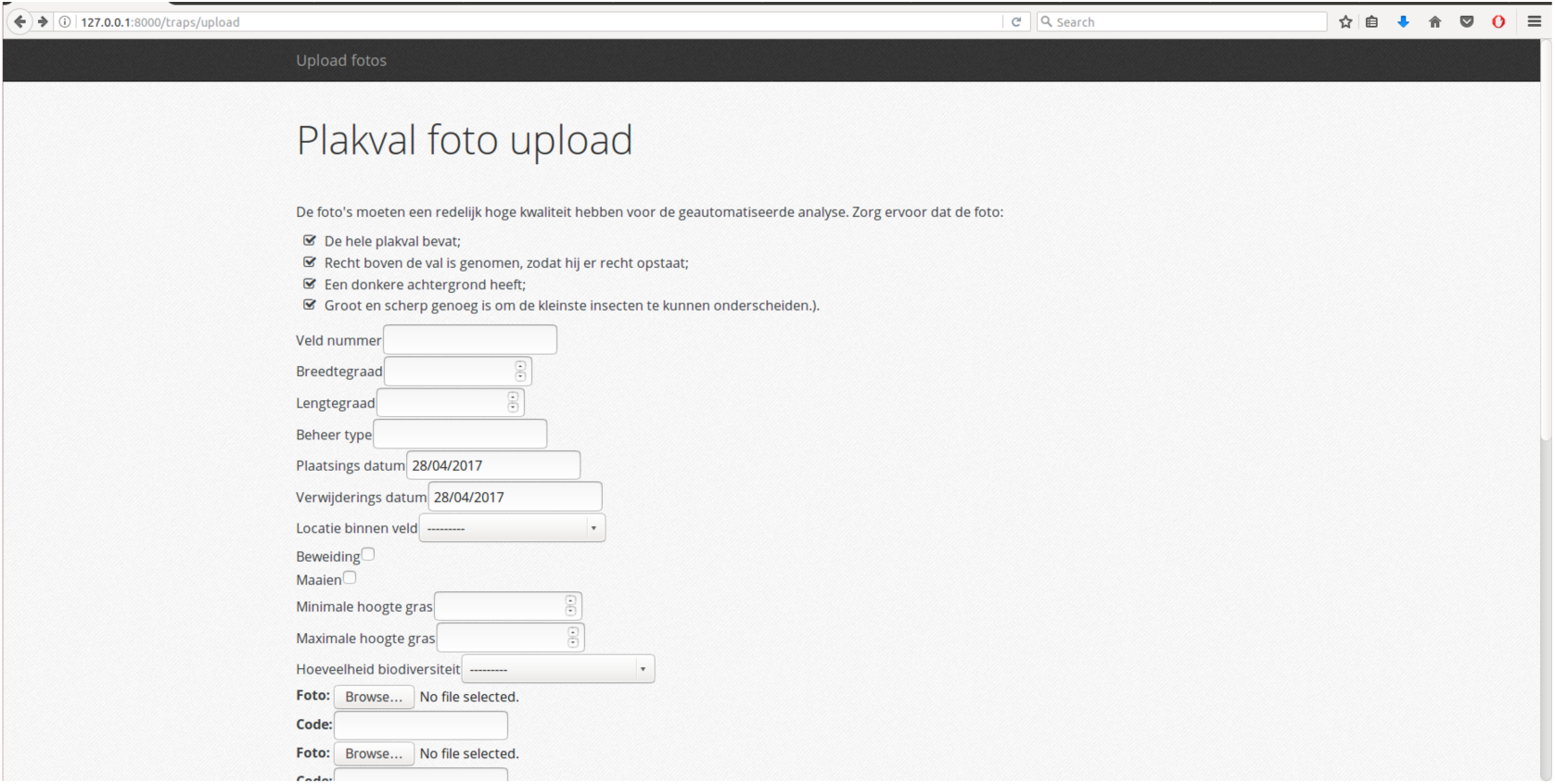
Website developed during the project. Visible is the uploads page where the information of the field can be entered. At the bottom of the image the first field dedicated for the uploading of images is visible

The website developed during the project also has several steps of the workflow.

1. The visitors are greeted with a homepage with information on the larger project, and a reminder of the requirements of the photos. There is a link redirecting them to the upload page.
2. The upload page then asks them to submit relevant information. This includes the name of the farm where the field is located, the geographic location using the google maps API, the estimated range of the height of the grass, and a rough estimate of the species richness in the field. On the same page, the user is asked to upload the images they took of the traps deployed in the field.
3. After submitting the photos and the relevant information the webpage calls the automated analysis script^15^ for each of the submitted photos. The results of the analysis are then automatically added to a file in a tab-delimited text file for analysis^16^, and the average of the areas is added together with the information gathered in step 2 to a second text file^17^.
4. The user is redirected to a different page where the results for this field are presented, and a link is presented to step 2 to keep little time between the uploading of different forms.

### Fieldwork

The traps used were 10cm × 24.7cm in diameter, with a uniform yellow colour and a standard size to aid in the automated analysis. The standard dimensions are used to automatically determine the size and scale of the photos and to be able to give accurate results. The colour yellow was chosen to lure the insects towards the traps in order to ensure high enough insect densities on the traps. No pre-printed lines were present on the traps to avoid false positives in the script, and to maintain accurate results.

The traps were deployed during two rounds of fieldwork in May. During both rounds of fieldwork, the traps were set within a few hours of each other by volunteers and collected approximately 2 days after the traps were deployed. The first round of fieldwork started on the 1^st^ of May and the second round of fieldwork started on the 15^th^ of May. Traps were deployed in fields belonging to 6 different agricultural companies and these field were managed according to 4 different known management types. This large variation in management types was selected for this data to be used in a larger scale study on the status of the food supply for meadow bird chicks, and for the validation of this project to be more accurate and complete.

10 traps were deployed in each field during both rounds of fieldwork. These traps were positioned in a diagonal line across the field, with gaps of 10 meters between the traps, and a distance of at least 10 meters between the traps and the edge of the fields. See Fig 4. The traps were placed diagonally to capture the variation introduced by the presence of small ditches in the fields. All traps were deployed vertically around 10-15cm above the soil to avoid any debris landing on the trap. The grass surrounding the traps was trimmed to avoid any blades of grass touching the traps themselves.

**Fig 3.**
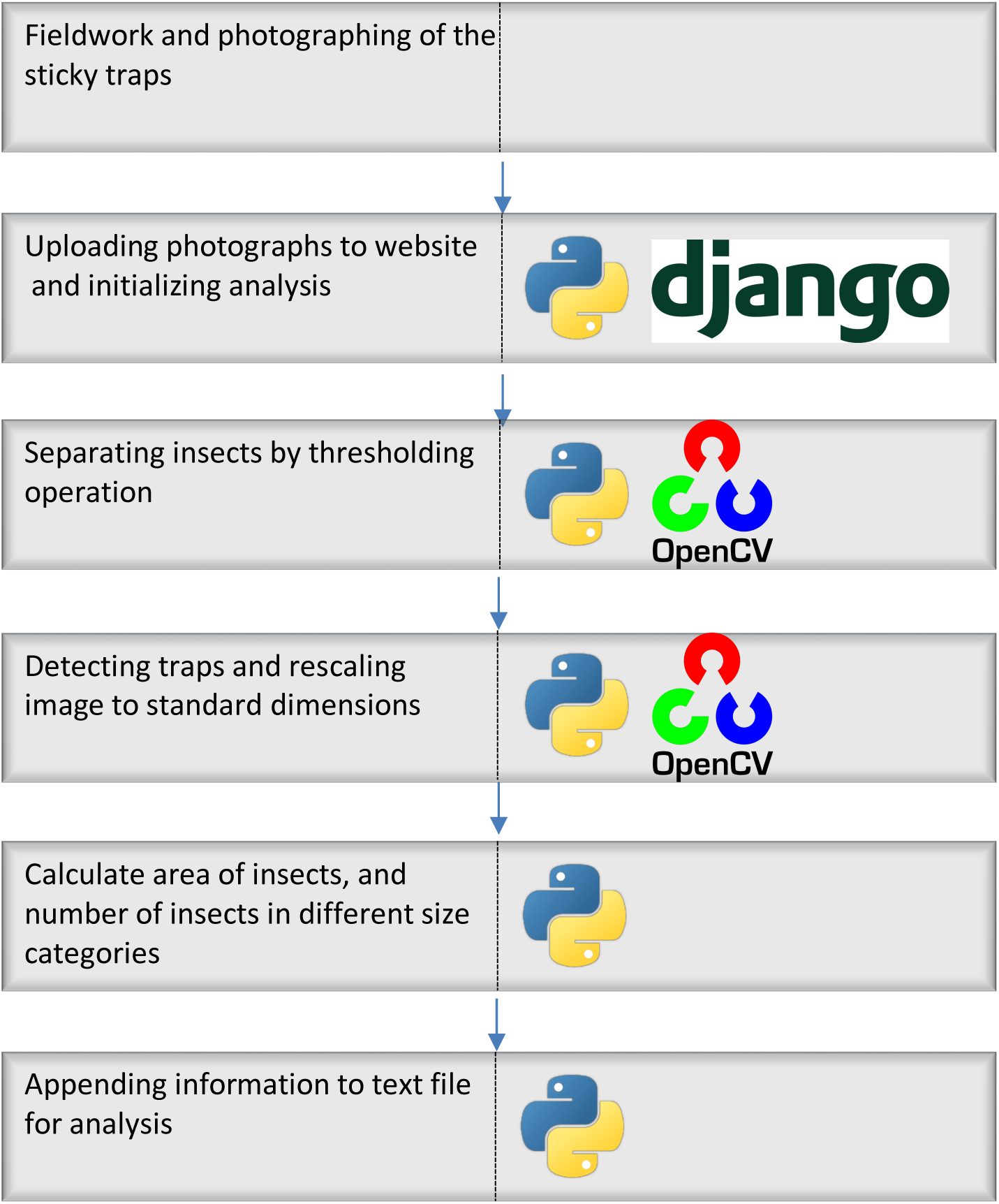
Workflow of the automated scripts. For each step the packages used are listed.

**Fig 4.**
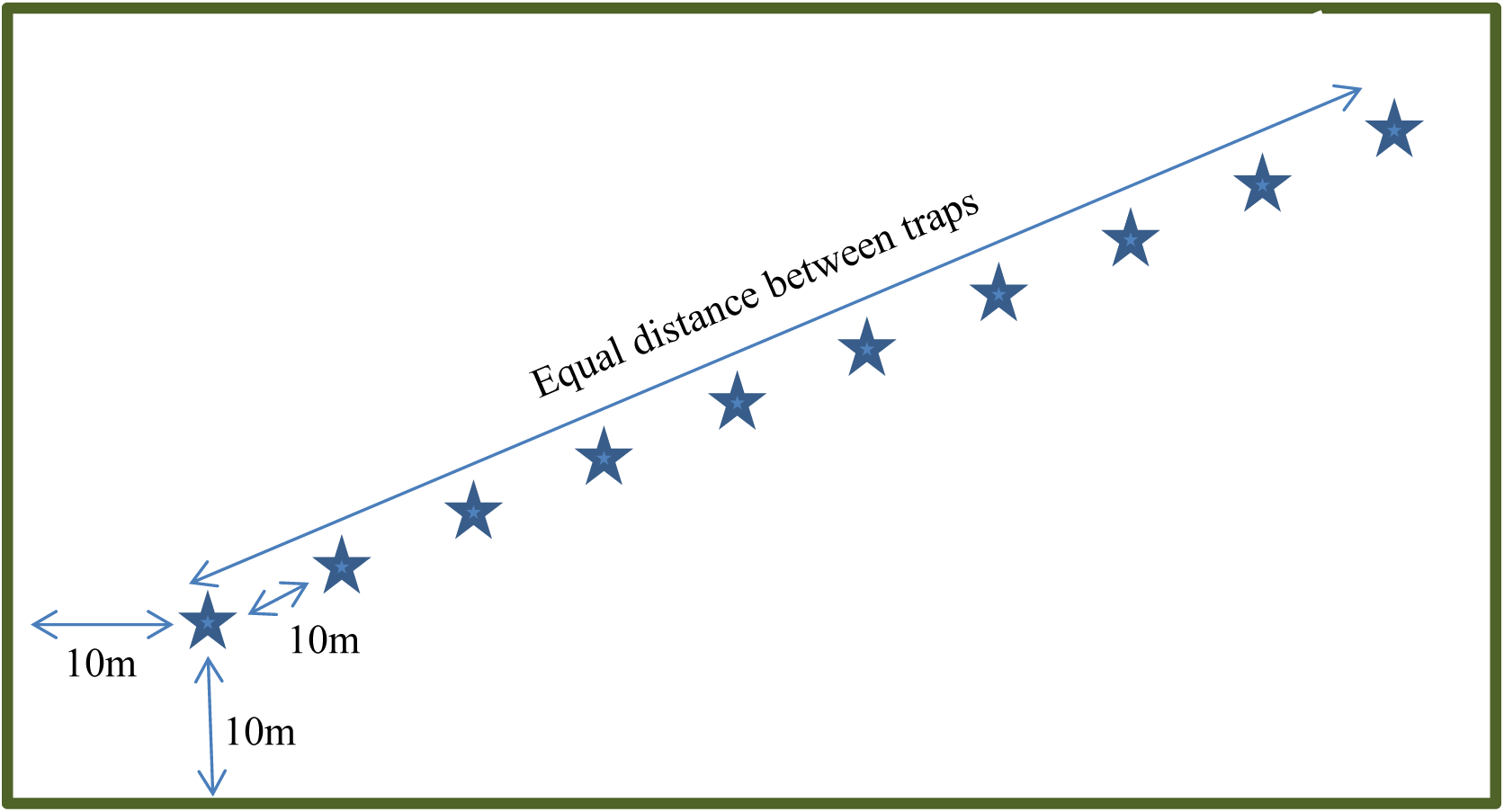
Deployment of the traps in the field. Each star represents 1 trap.

After collecting the traps, photographs were taken and submitted by the volunteers. The physical traps were also collected to count the number of insects caught on them. 110 traps were counted by hand. We also photographed all the counted traps using both my cell phone, and a high-end consumer camera. All photographs were analysed by the program, and all results were entered in one large datafile. This resulting data was then used for the analysis.

### Statistics

The hand counted data was converted to indicators representing the expected area and volume of the insects caught on the trap. These indicators were calculated using the number of insects of each size class caught on the trap and an estimate of the average size of insects in each size class. For each size class, the number of insects where multiplied by the square of the average length to obtain an indication for the area of the insects, and multiplied by the cube of the average length to obtain an indication of the volume present. The values for the different size classes are added to give one Correlations with the output of the program were performed with the indicators for the area and volume. The number of insects in each size class as calculated by the program were correlated with the actual counts. To verify that the patterns in the data returned by the script are the same patterns in the hand counted data a boxplot was made where the data was split by date, company, and management practice.

## Results

All correlations investigated were highly significant, with p-values <0.001. This is needed for a predictive program, but we are more interested in the remaining variance. We have therefore focused the results on the R^2^ of the correlation which gives an indication of the variance explained by the correlation.

Plotting the results of the program against the indicator for the area gives a good correlation, with an R^2^ of 0.72. (Fig 5a). There appears to be no effect of the different cameras, dates, companies, and management practices on the regression line drawn (Fig 5b-d). We do see that there is slightly more variation in the photographs taken by the volunteers (Fig 5b). This is most likely because of differing lighting conditions and photo qualities. The photographs taken by the phone camera and the high-end consumer camera show the same amount of variance. This indicates that as long as the resolution of the photographs is sufficiently high, and the photographs are taken in a standardized fashion, then the actual camera used to take the photograph is not of influence on the results obtained by the script.

**Fig 5a.**
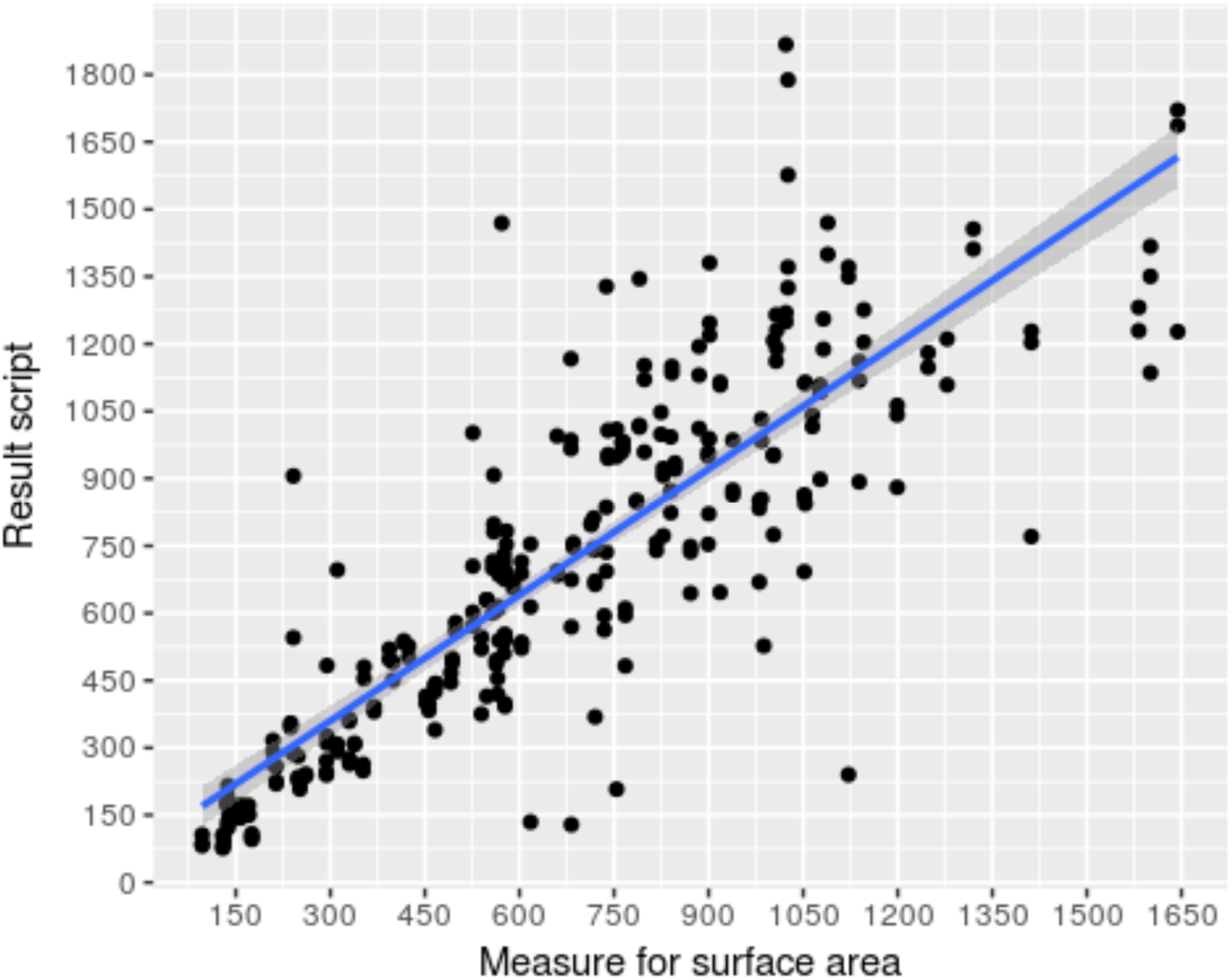
The ouput given by the program on the y axis plotted against the measure of the surface area as calculated in the methods, with regression line.

**Fig 5b.**
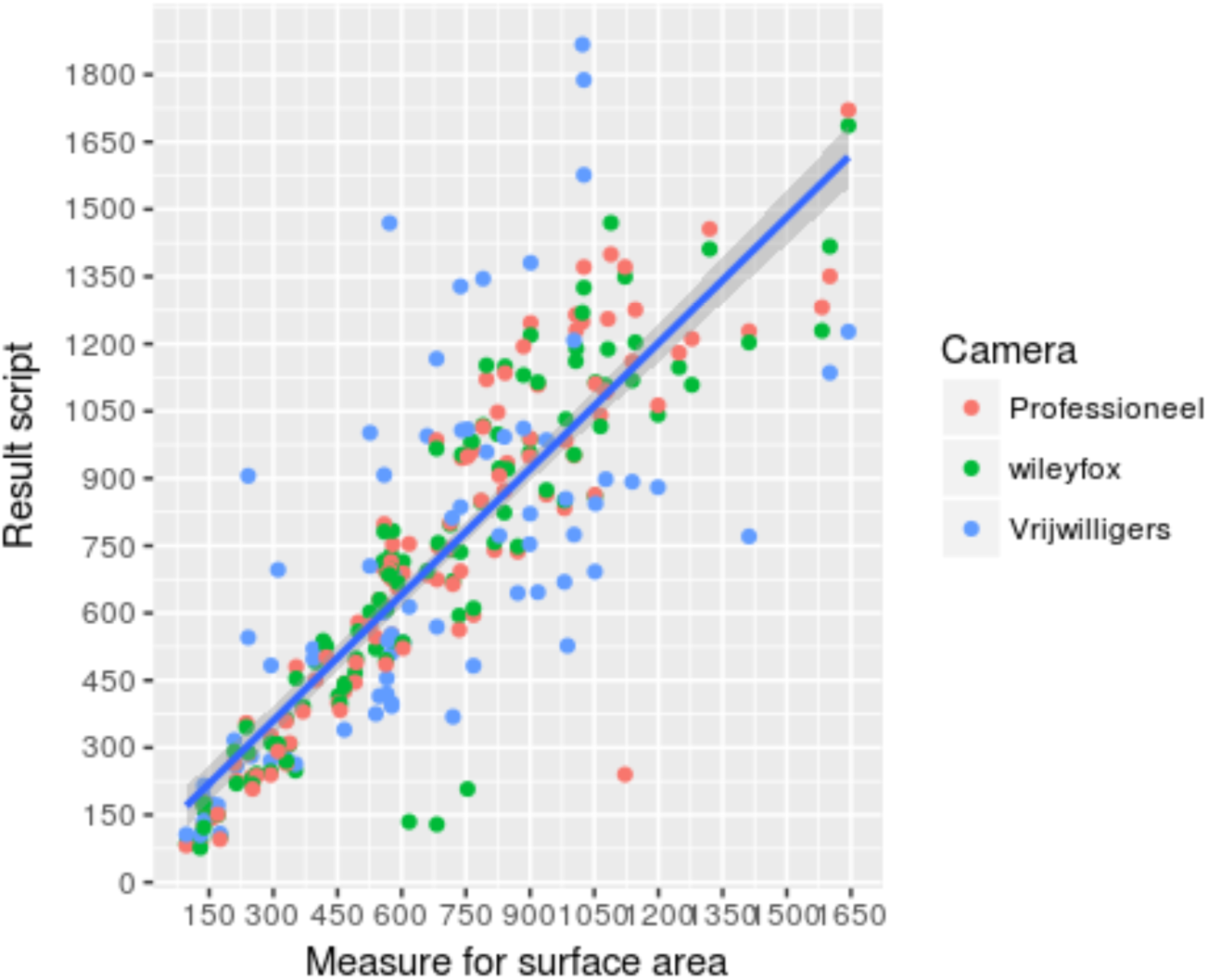
the output of the program plotted against the counted surface area. The points in the plot are coloured according to the camera the image was taken with. The photos taken with the professional camera and the camera phone seem to have less variance; no other patterns emerge influencing the regression. We can however conclude that using a standardized method of taking images will result in more accurate data.

**Fig 5c.**
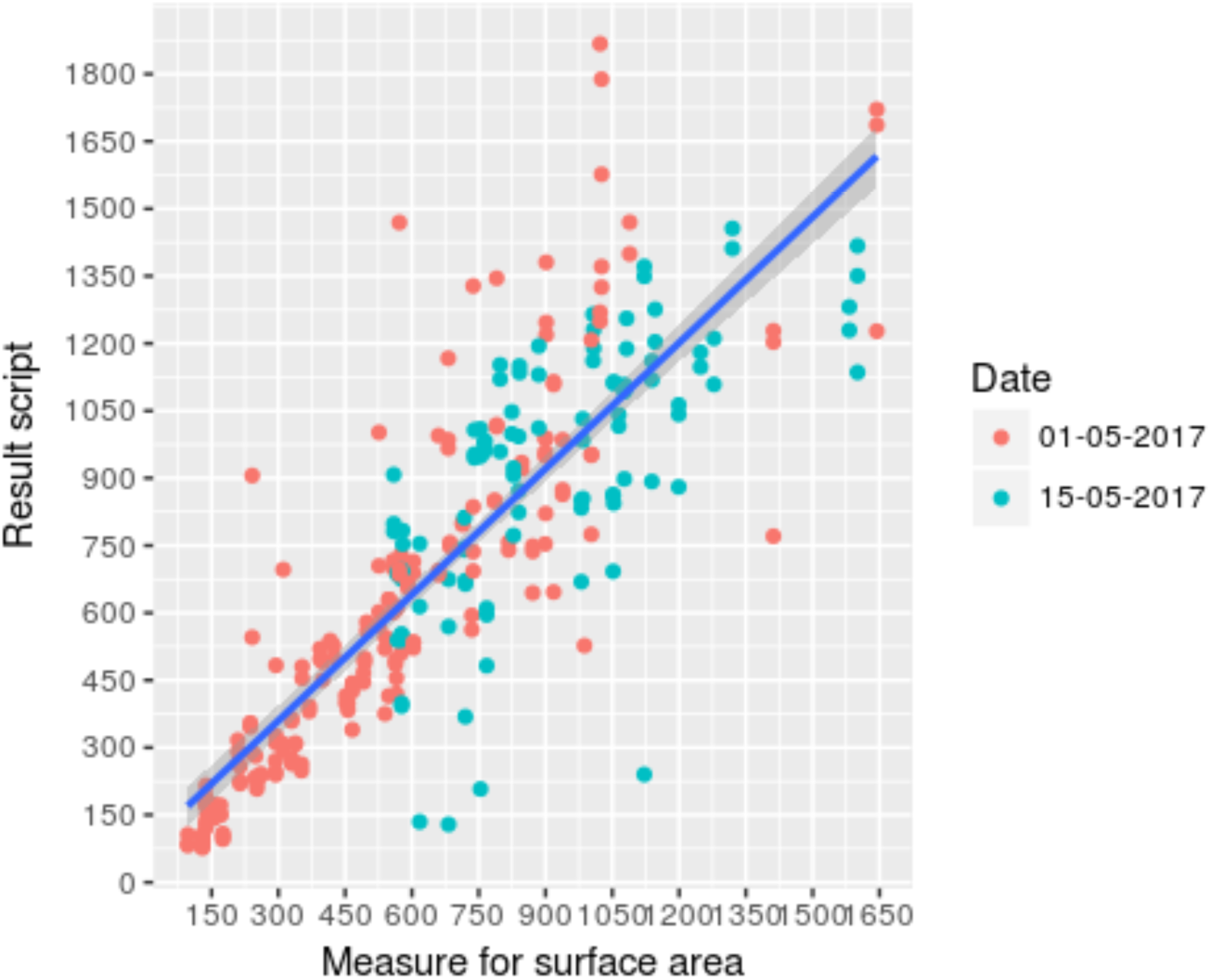
the output of the program plotted against the counted surface area coloured by date. We see that there is considerable clustering present. Most data points of the first round of fieldwork can be found nearer the origin. This is expected because there are fewer insects present in the fields during the first round of fieldwork.

**Fig 5d.**
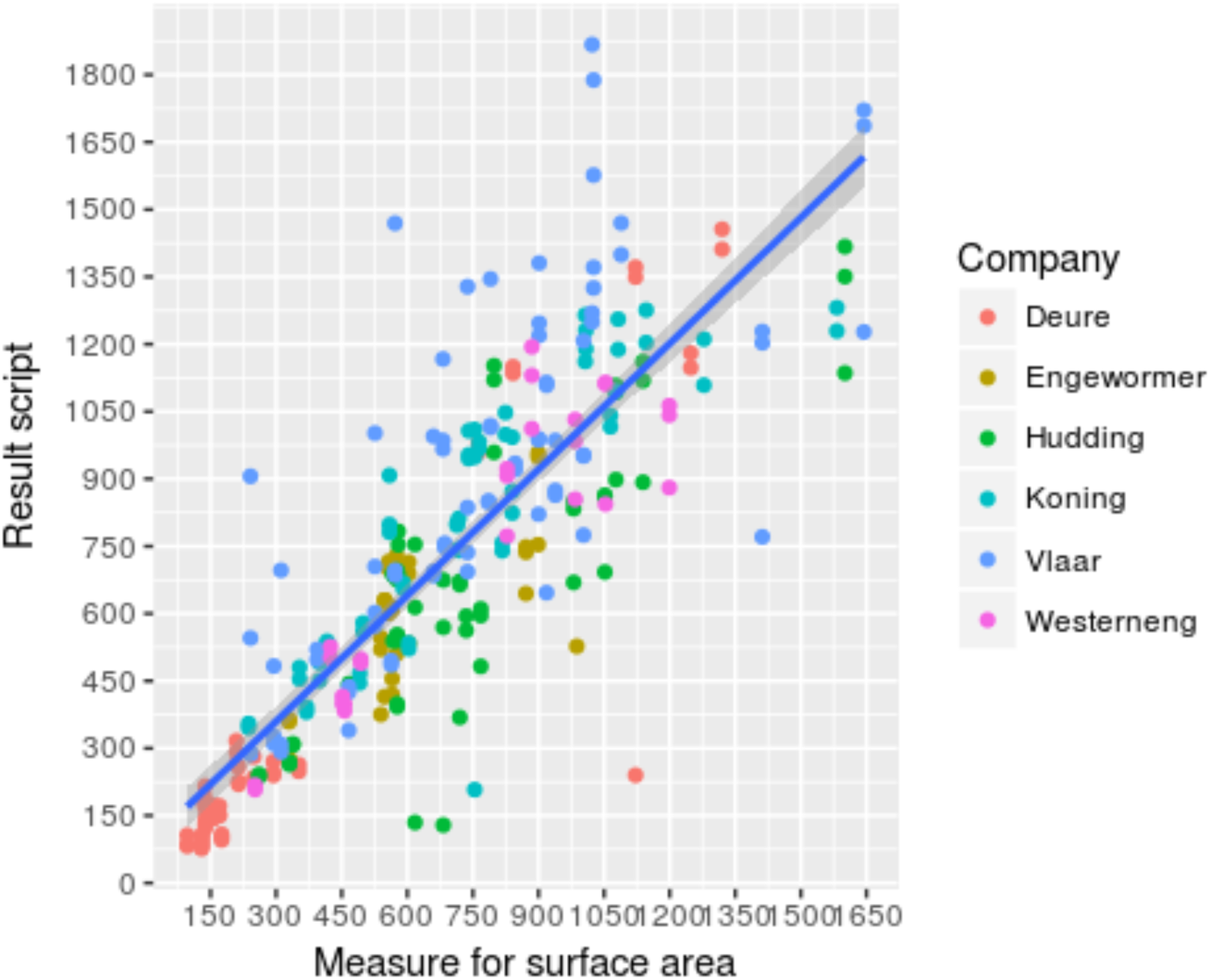
The output of the program plotted against the counted surface area coloured by the agricultural company at wich the trap was set. We find some clustering in the data points. This is also expected since we expect that traps frome the same field, and by extension the same company, will have similar numbers of insects present.

We also see a strong clustering of the different dates in the various plot. (Fig 5c) This can be explained as an effect of the date itself, as there were more insects present on the traps set on the 15^th^ of May. This clustering however does not have a noticeable effect on the regression line obtained since the results of both dates are spread equally on both sides of the regression line. Based on these observations we can conclude that the script appears to respond solely to the insects present on the trap, as indicated by the absence of any pathological patterns in the different plots we have made.

The correlation between the surface area returned by the script and the manually determined volume of the insects on the trap was also strong, but slightly worse than the correlation with the manually calculated surface area of the insects present on the trap (Fig 5e) (R^2^ = 0.64). We see the same patterns emerging in the plots concerning the distribution of the data when coloured according to date, management type, and company. (Fig 5f – 4h).

**Fig 5e:**
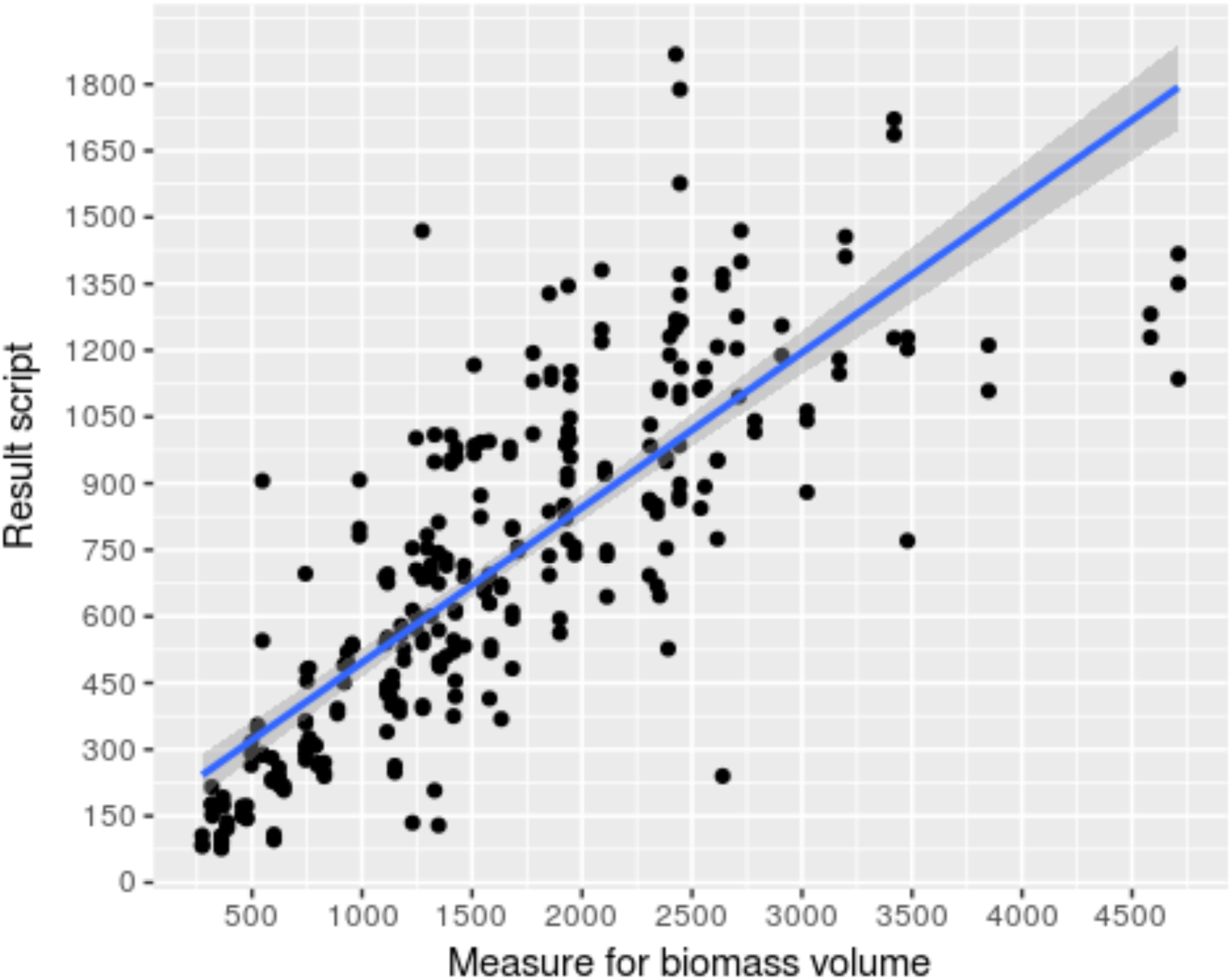
The total area returned by the script plotted against the hand counted measure of the volume of the insects present on the trap as calculated in the material and methods; with regression line.

**Fig 5f:**
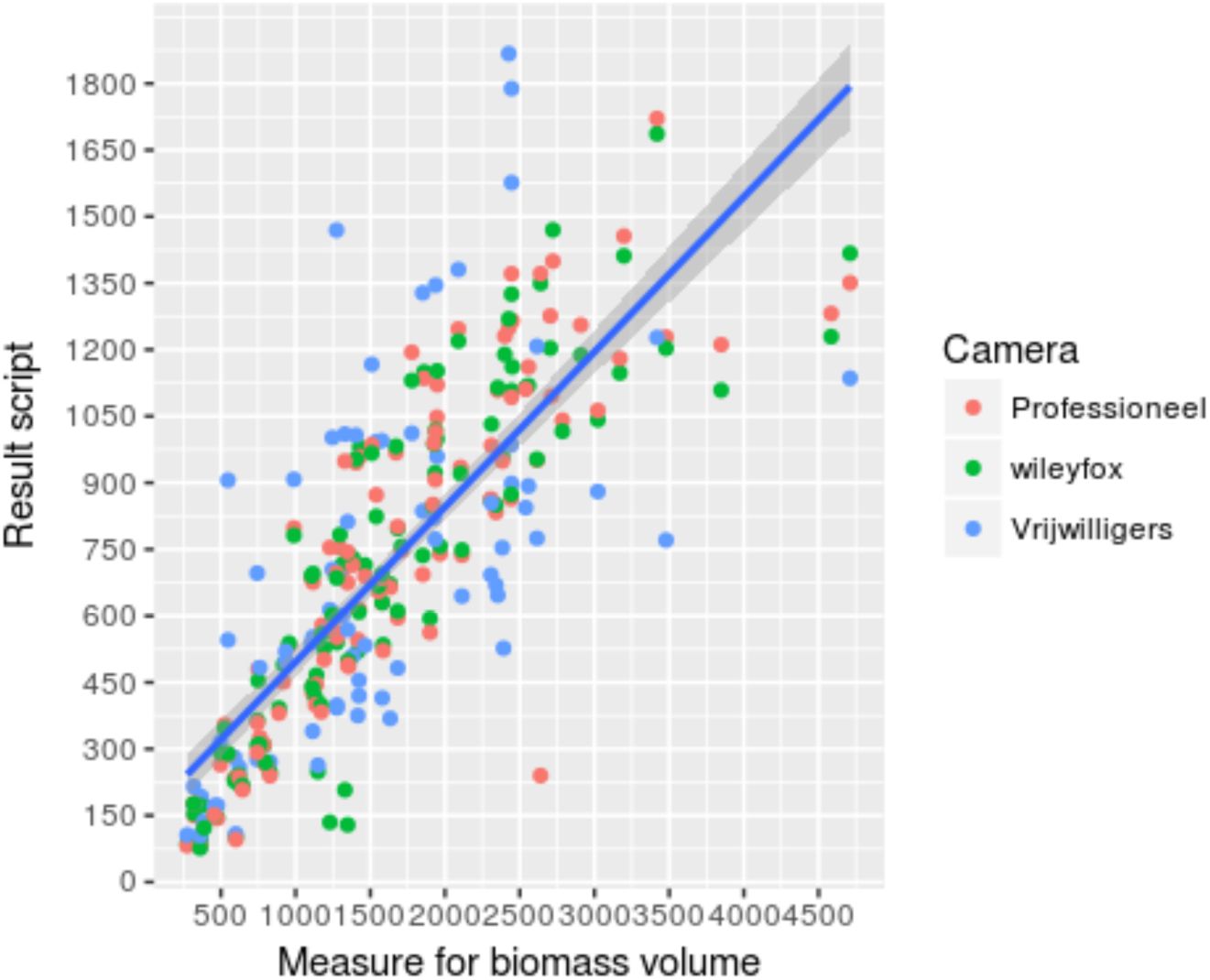
The regression of the area returned by the script and the estimated volume of the insects coloured by the camera the images were taken with. We see the same patterns in the plot of the estimated area: There is no apparrant pattern influencing the slope of the regression line, although the variance in the photographs taken by the volunteers is higher than the photographs taken by me.

**Fig 5g:**
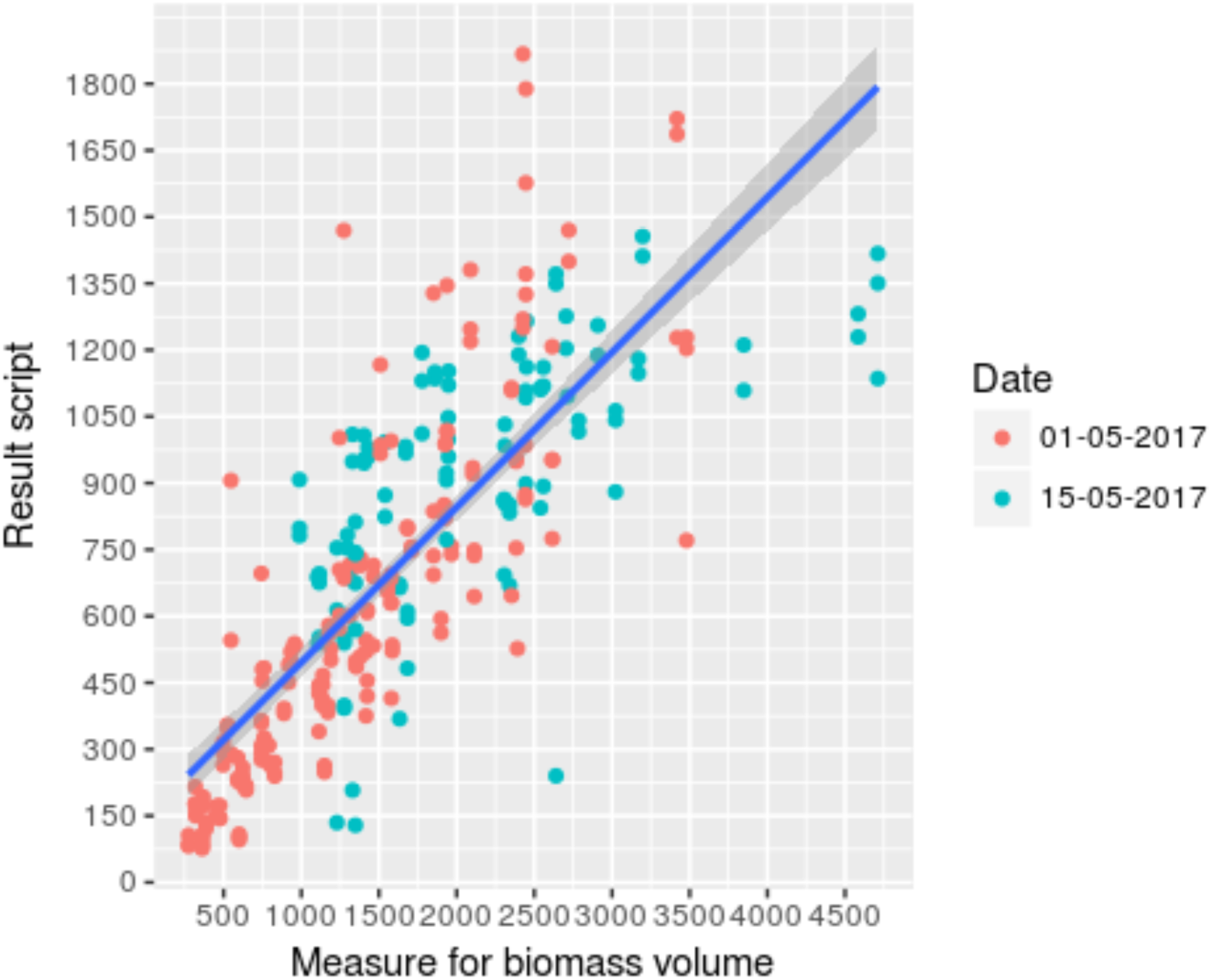
the area returned by the script plotted against the estimated insect volume coloured by date. Although there is significant clustering in the dates, this is expected and does not influence the slope of the regression line.

**Fig 5h:**
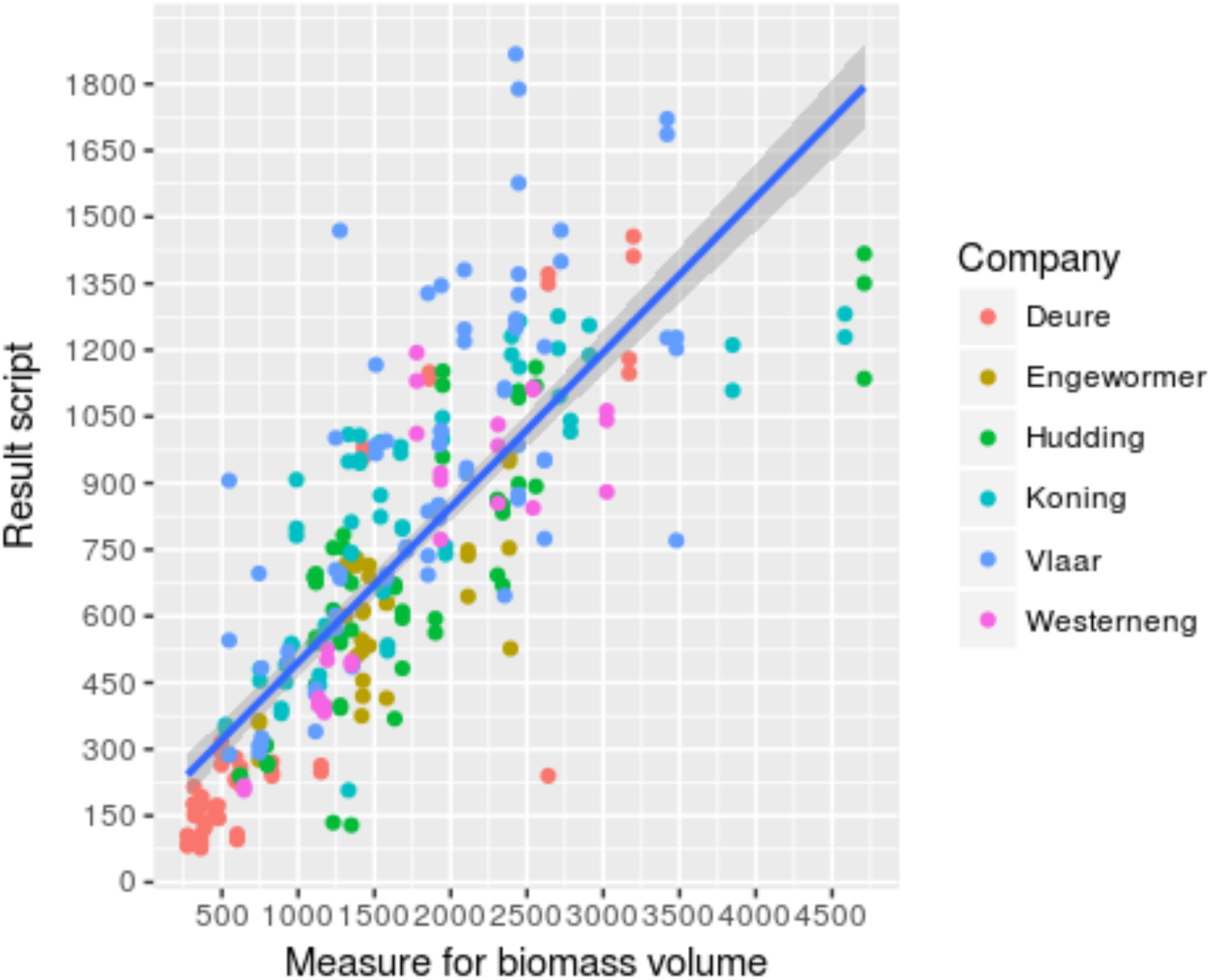
the area returned by the script plotted against the estimated insect volume on the trap coloured by the agricultural company where the traps were set. We can observe some clustering of the different companies. This clustering does not have an influence on the slope of the regression line.

The script appears to also give a good indication of the number of insects caught on the traps. The R^2^ of both the category 4mm and smaller (0.779) and the category between 4 and 10 mm (0.687) are comparable to the R^2^ of the area measurements. The amount of insect larger than 10 mm can however not be accurately predicted (R^2^=0.205) (Fig 6). This is because the smaller insects can overlap at higher densities and result in an overestimation of the number of these large insects as there is no way yet of distinguishing between one large insect and several small overlapping insects. The larger insects might also be slightly smaller in surface area than the threshold set in the script because of their shape. In this case this will lead to an underestimation of the number of large insects present. Combining these two effects results in an unreliable result for the number of large insects.

**Fig 6a.**
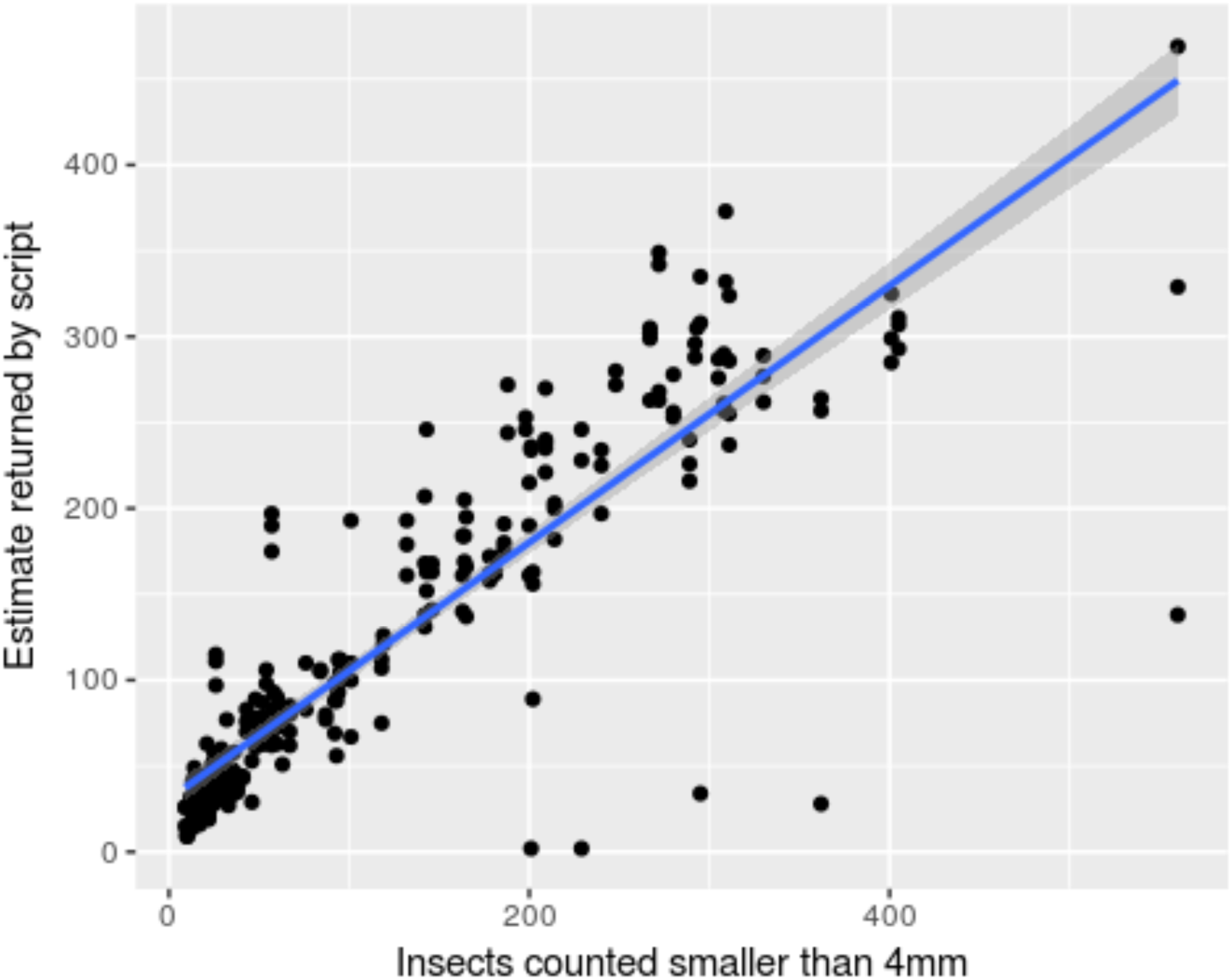
The number of insects smaller than 4 millimetres found by the script is plotted on the y axis against the number of insects smaller than 4 millimetres counted by hand. The regression line has also been drawn. There is only a small amount of residual variance in this regression, although there appear to be several outliers below the main point cloud. We can observe some clustering of the points, with the gap between these point clouds at 100 to 120 hand counted insects. The clustering however does not have an influence on the regressionline.

**Fig 6b.**
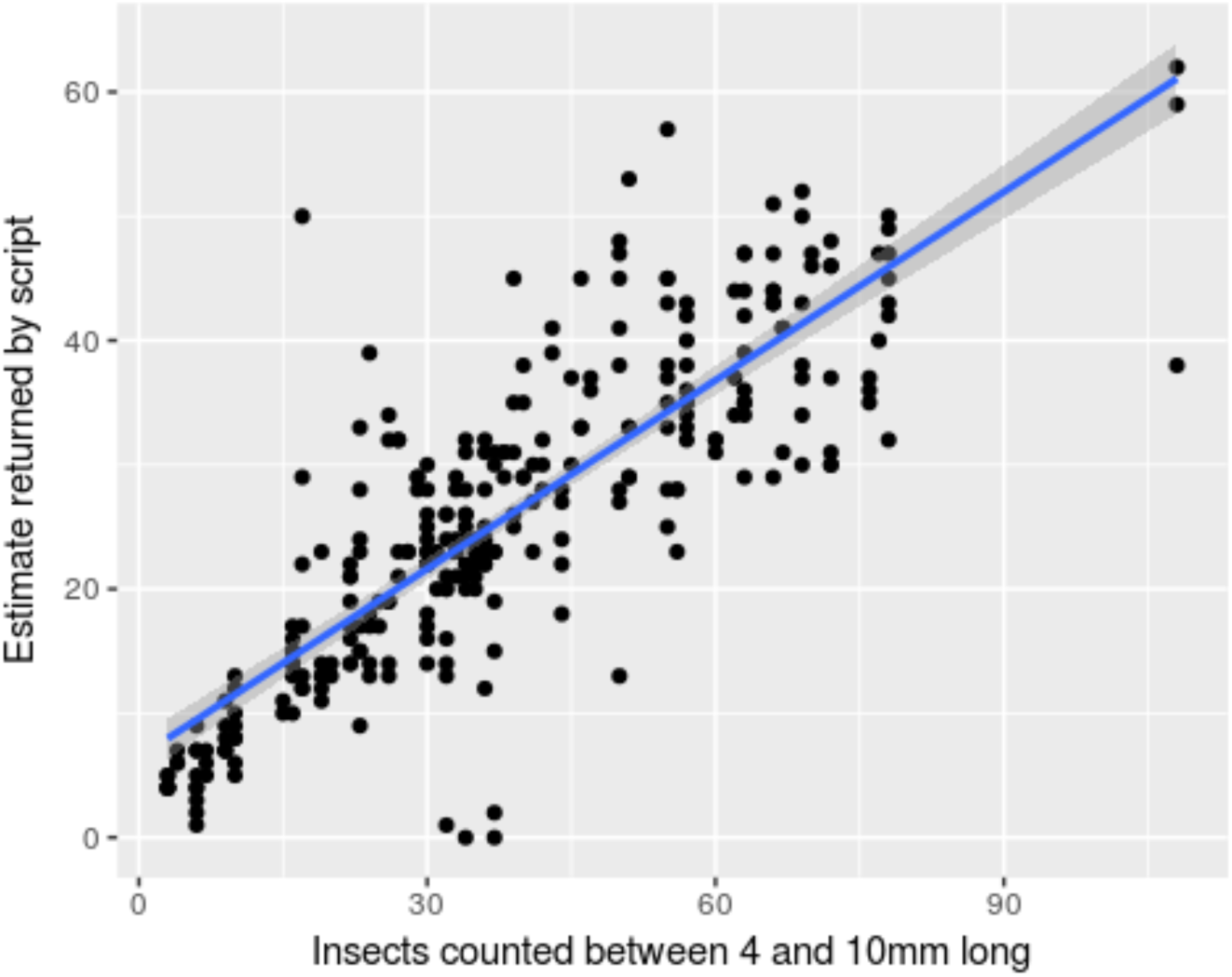
Insects estimated to be between 4 and 10mm by script on the y axis plotted against insects hand counted between 4 and 10mm. The regression line has also been drawn. There is a significant amount of residual variance in this regression, and there appear to be several outliers below the point cloud as well. We can not see any clustering of the points.

**Fig 6c.**
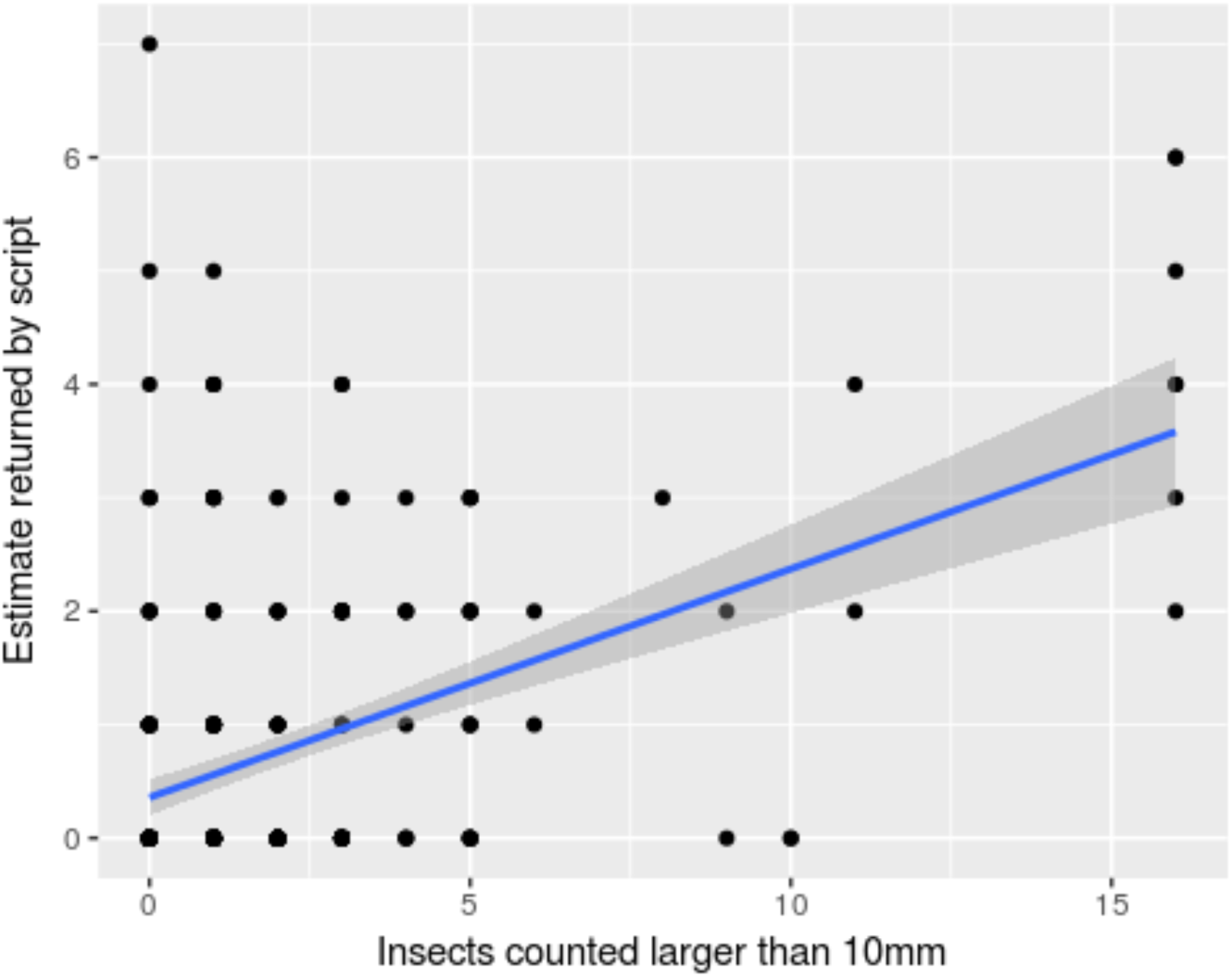
Number of insects larger than 10 mm estimated by script on the y axis plotted against insects hand counted larger than 10mm. The regression line has been drawn because the correlation was significant. The amount of residual variance in this plot means that this estimate is not useful in the estimation of the biomass since the regression only explains a small amount of the variation in the returned values of the script.

The patterns we find in the data are consistent. This is visualized using boxplots in Fig 7. We see that the same patterns emerge in both boxplots, with roughly equal variance. This indicates that the use of this program does not significantly influence the results gained from the data.

**Fig 7a.**
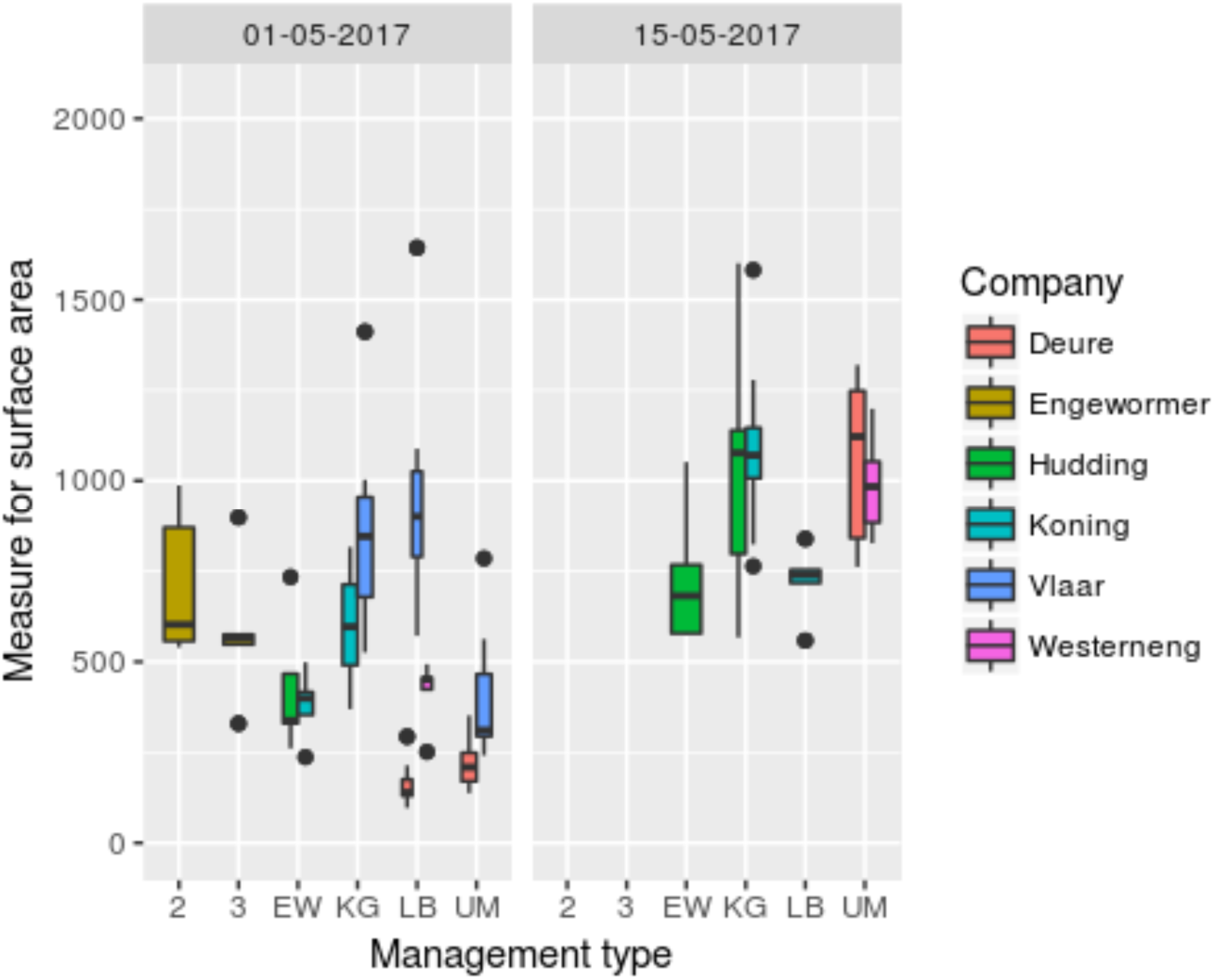
The estimate of the manually estimated insect surface area is plotted on the y axis, and plotted by date, agricultural management type and company. We find a relatively large amount of variance within different fields.

**Fig 7b.**
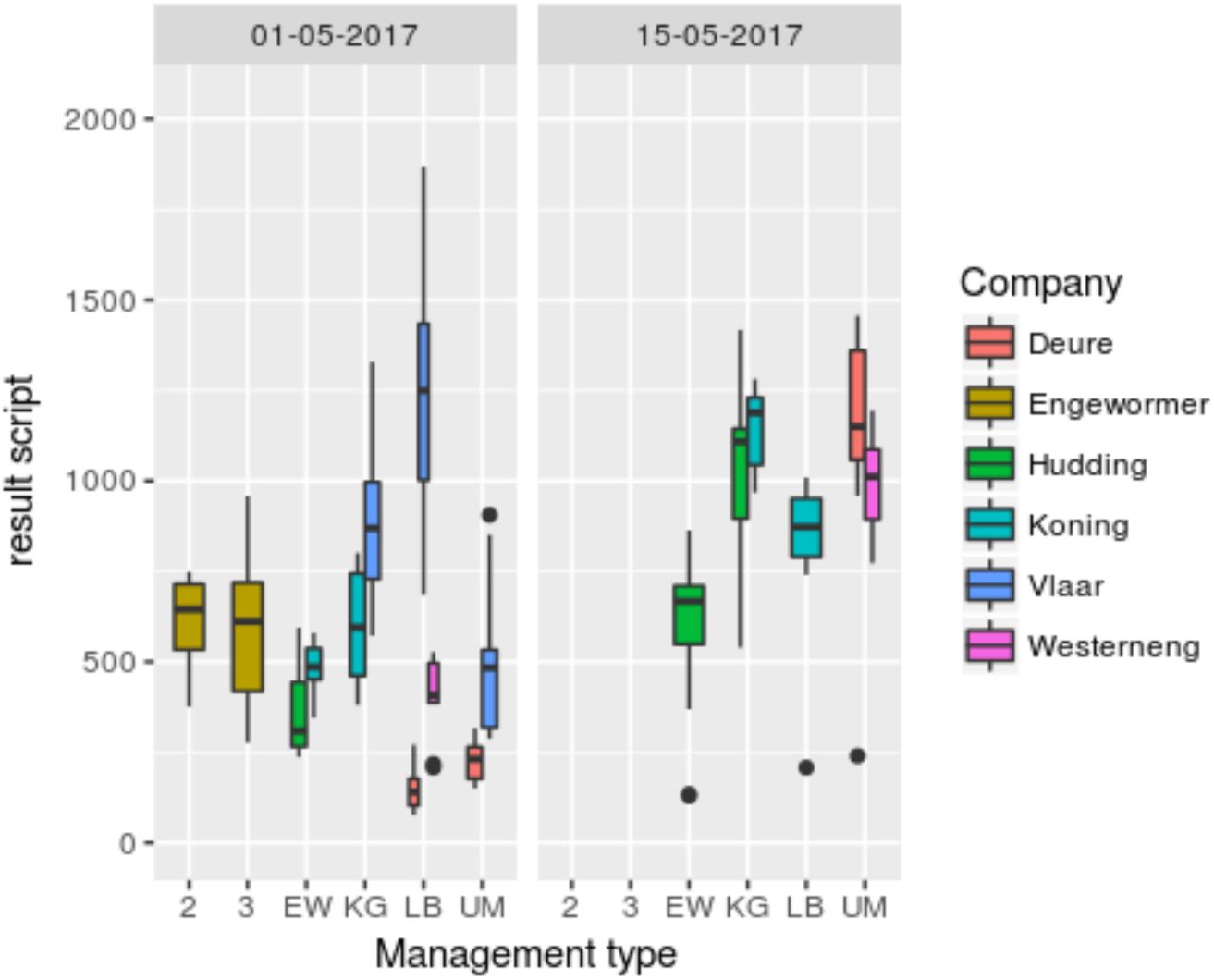
The area returned by the script plotted on the y axis according to the date, agricultural management policy and agricultural company where the traps were set. We find the same patterns in the plot of the manual estimate of the area. This indicates that the results of the script do not have an influence on the results and the conclusions that can be drawn from the data. The relative variance of the result of the script is equal to the variance found in the estimates made by hand. This indicates that the residual variance in the results of the script does not have an influence on the total variance when compared to the manually gathered data

## Discussion

Although the variance in the regressions is higher than perhaps desirable, we do find that the patterns found in the box plots of the hand counted data and the data obtained by the script are very similar. In fact, all but one of the medians of the hand counted data fall between the 1^st^ and 3^rd^ quantiles of the data after the output of the script is rescaled according to the results of the regression. This would indicate that this script can give an indication of the variance present in different areas and across different management types. None of the correlation coefficients found in the regressions were 1. The residual variance however is always relatively low. This means that the program usually over- or underestimates a certain parameter by a set factor. In the case of the area returned by the script correlated with the manually determined area the area calculated by the script was consistently 7.5 times higher than the manually derived value. This is a result of the way the indicator for the area of the insects was calculated. This is a result of the way the manual area was calculated. The relative increase in area was used instead of the real values to increase the simplicity of the calculation, so that the values did not represent the actual areas but instead the ratios between the different size classes. Because the smallest size category had a relative increase in area of 1, this meant that all values in the manually counted data should be multiplied by ∼2^2^ or ∼3^2^ eliminating this anomaly.

When investigating the regression plots closely, we observe that the results near zero are further below the regression line. The resulting pattern resembles the pattern observed from a function containing a square root. This effect is not very pronounced however. When trying to use a log-log plot to make the data entirely linear, we find a higher amount of variance, with several very large outliers in the data. Because the effect is only slight we have elected to use a linear model instead. We believe that this effect arises as a result of the normal operation of the script. Because the script uses an adaptable threshold when finding the insects, it could be that the script is more sensitive to insects at low insect densities, meaning that some noise might be seen as an insect. Future research will have to determine if this noise can be eliminated. We expect that setting a lower bound on the size of insects will help in eliminating this pattern, however we have not been able to implement this yet.

We find that the number of insects larger than 10 mm cannot be accurately predicted. We believe that this is because of 2 factors. The first is that there can be a considerable overlap in the insects counted on the more densely covered traps resulting in an overestimation of the number of large insects present on those traps. Secondly, the shape of most, if not all, insects larger than 10 mm is not round, but instead the width is considerably shorter than the length of the insect. This will most likely result in the area of the large insects being smaller than the threshold value that we have set for the category. This in turn will result in an underestimation of the number of large insects present on the trap.

Not being able to find the largest insects however is not crucial to the working of the program, as the main food source for the meadow bird chicks is the insects between 4 and 10 mm in size. The biggest insects are still a good source of food, but they are usually quite rare when compared to the number of smaller insects.

The number of insects present on the traps is underestimated for both other size categories. The slope of the regression line in the category of 4 mm and smaller is 0.75, which means that a quarter of the insects is either missed, or is classified as a larger insect as a result of overlapping with other insects. This slope is 0.5 for the insects in the size category between 4 and 10 mm. This is still a constant result, so an approximate prediction can still be made. We believe that there are most 3 likely reasons for this underestimation.

1. The insects that are only slightly larger than 4 mm are most likely misclassified as being less than 4 mm. This is because insects do not have a perfectly square shape, and are in fact oblong.
2. Some insects might be too far degraded to be recognized as one insect, therefore also being misclassified as less than 4 mm. This will be rare however as most insects are reasonably intact.
3. Finally, there will be some insects that are overlapping with another insect and are then categorized as being more than 10 mm.

## Conclusion

We have developed a program that is able to return an estimate of the insect biomass present on a sticky trap from an image of the trap. The script in its current state does allow for considerable improvement and optimization in future. This includes, but is not limited to:

- making the algorithms for finding the traps and the insects more robust and accurate;
- decreasing the computational time needed and lowering the memory required;
- improving the look of the website; and
- optimizing the output for analysis.

The application of this script is not just limited to meadow birds: several researchers have shown interest in using this script in assessing the food availability for other groups of birds, or animals in general. The only constraint here is that the traps must be set in the same location that the animals forage for insects, as well as using traps suitable for this script: namely yellow traps without writing or lines present.

Future projects might involve the identification of the insects caught on the traps. This will almost certainly not be to the species level; however, it might be possible to identify the family or genus of certain insects caught on the traps using one or more techniques such as neural networks. Software to identify pests caught on sticky traps in greenhouses has already been developed. This software however only needs to identify a small number of different species, and the identification is therefore a lot simpler when compared to the vast number of different insects found in fields, especially when the floral diversity is higher.

## Footnotes

1 https://github.com/naturalis/imgpheno

2 http://opencv.org/

3 https://www.djangoproject.com/

4 https://developers.google.com/maps/

5 https://pypi.python.org/pypi/django-geoposition

6 https://github.com/naturalis/imgpheno/blob/master/examples/sticky-traps.py#L146-L156

7 https://github.com/naturalis/imgpheno/blob/master/examples/sticky-traps.py#L83

8 https://github.com/naturalis/imgpheno/blob/master/examples/sticky-traps.py#L160-L168

9 https://github.com/naturalis/imgpheno/blob/master/imgpheno/__init__.py#L865-L877

10 https://github.com/naturalis/imgpheno/blob/master/imgpheno/__init__.py#L845-L862

11 https://github.com/naturalis/imgpheno/blob/master/examples/sticky-traps.py#L114-L116

12 https://github.com/naturalis/imgpheno/blob/master/examples/sticky-traps.py#L172-L179

13 https://github.com/naturalis/nbclassify/blob/sticky-traps/html/sticky_traps/analyze.py#L111-L139

14 https://github.com/naturalis/nbclassify/blob/sticky-traps/html/sticky_traps/results/foto_data.txt

15 https://github.com/naturalis/nbclassify/blob/sticky-traps/html/sticky_traps/analyze.py

16 https://github.com/naturalis/nbclassify/blob/sticky-traps/html/sticky_traps/results/foto_data.txt

17 https://github.com/naturalis/nbclassify/blob/sticky-traps/html/sticky_traps/results/veld_data.txt

## References

Barbedo, J. G. A. (2014). Using digital image processing for counting whiteflies on soybean leaves. Journal of Asia-Pacific Entomology, 17(4), 685–694. https://doi.org/10.1016/j.aspen.2014.06.014

Beintema, A. J., Moedt, O., Ellinger, D., Beintema, A., & Moedt, O. (2007). Ecologische Atlas van de Nederlandse weidevogels. Haarlem: Schuyt & Co. Retrieved from http://www.beintema.com/atlas.html

Deru, J., Noordijk, J. (kenniscentrum E., Luske, B., & Wennekens, E. (2016). Meten van voedsel voor weidevogelkuikens. Tussen Duin En Dijk, (1), 3. Retrieved from http://www.louisbolk.org/nl/publicaties/publicatie/?pubID=3118

Noordijk, J. (2014). Meten van weidevogelvoedselbeschikbaarheid door vrijwilligers. Kenniscentrum EIS

Schekkerman, Hans; Boele, A. (2008). Precocial problems. Shorebird chick performance in relation to weather, farming, and predation. PhD, University of Wageningen. Retrieved from http://library.wur.nl.ezproxy.leidenuniv.nl:2048/WebQuery/wurpubs/wever/366754

Solis-Sánchez, L. O., García-Escalante, J. J., Castañeda-Miranda, R., Torres-Pacheco, I., & Guevara-González, R. (2009). Machine vision algorithm for whiteflies (bemisia tabaci Genn.) scouting under greenhouse environment. Journal of Applied Entomology, 133(7), 546–552. https://doi.org/10.1111/j.1439-0418.2009.01400.x

Sovon. (n.d.). Grutto | Sovon.nl. Retrieved March 6, 2017, from https://www.sovon.nl/nl/soort/5320

Pereira, S., Gravendeel, B., Wijntjes, P., & Vos, R. (2016). OrchID: A Generalized Framework for Taxonomic Classification of Images Using Evolved Artificial Neural Networks. bioRxiv, 070904.

